# Exploring The Genome of The Oribatid Mite, *Oppia Nitens* – Environmental Stress Response and Toxicity Adaptation

**DOI:** 10.1101/2024.10.27.619232

**Authors:** Adedamola A. Adedokun, Hamzat O. Fajana, Olukayode O. Jegede, Austin S. Hammond, Derek D.N. Smith, Stephanie Kvas, Thulani Hewavithana, Lingling Jin, Juliska Princz, Steven D. Siciliano

**Author notes:** Corresponding author: Department of Soil Science, University of Saskatchewan, Saskatoon, SK 11 S7N 5A8, Canada. Current address: Global Institute for Food Security, University of Saskatchewan, Saskatoon, SK S7N 4J8, Canada.

## Abstract

Oribatid mites are one of the most abundant groups of microarthropods in soil. *Oppia nitens*, belonging to the family Oppiidae, one of the largest and most diverse families of oribatid mites, has been developed as a standardized model test organism for assessing soil contamination. However, the limited availability of genomic information for this species hinders our understanding of its physiological adaptation and sensitivity to chemical and environmental stressors in soil. Hence, we present the annotated *O. nitens* draft genome assembled using both Oxford Nanopore Technologies and Illumina sequencing platforms as a basis to identify potential genes that can be linked to adaptation to chemical and environmental stressors.

The sequences were assembled into 65 scaffolds spanning 125.4Mb with a 24.5% GC content and an N50 length of 4.41Mb. Genome quality and completeness were checked using arthropod Benchmarking Universal Single-Copy Orthologs (BUSCO) analysis, which identified 93.5 % complete single-copy orthologs, 3.4% complete but duplicated orthologs, 0.5% fragmented, and 2.6% missing orthologs (n=2934). The NCBI Eukaryotic Genome Annotation Pipeline annotated 15,291 genes, 16,969 mRNAs, and 14,938 proteins.

Here, we describe the *O. nitens* complete draft genome and discuss its utility as a genetic basis for further investigations and understanding of the molecular mechanisms and physiological functions in adaptations to environmental change, especially tolerance to metal stress.

## 1. Introduction

The emergence of (eco)toxicogenomics has expanded the scope of soil toxicology and risk assessment beyond traditional approaches that focused on apical endpoints related to survival and growth. By using genomic methods, researchers can gain insight into the mechanistic basis of biological effects, moving from whole-organism outcomes to molecular events and adverse outcome pathways (van Straalen and Feder, 2012). The application of genomic and transcriptomic tools for soil organisms has reached significant milestones for elucidating chemical components of toxicological significance within environmental samples (e.g., Nota et al., 2009; Chen et al., 2014, Zhu et al., 2020; Lin et al., 2021). However, while classical soil ecotoxicology has established standardized species and test methods that represent diverse ecological niches and trophic levels, toxicogenomic profiles of certain standardized species are still lacking, hindering our ability to fully comprehend the linkages between soil pollution and organism and ecosystem health.

Toxicogenomic profiles enhance our understanding of mechanisms of action by systematically revealing how environmental stressors influence gene expression, disrupt cellular pathways, and trigger molecular responses, thereby providing detailed insights into the biological impacts of toxic exposures (Brinke and Buchinger, 2017). Toxicogenomic profiles exist for some soil species including *Folsomia candida*, *Sinella curviseta*, *Orchesella cincta*, and *Holacanthella duospinosa*, representing the collembolan model species (de Boer et al., 2009 and 2015; Faddeeva-Vakhrusheva et al., 2017; Zhang et al., 2019; Faddeeva-Vakhrusheva et al., 2016; Wu et al., 2017). These studies employed gene expression analysis to examine how springtails respond to environmental stressors and developmental processes. For instance, Lin et al. (2021) used transcriptomics and metabolomics to investigate the toxicity of antimony (Sb) to *F. candida*, finding that prolonged exposure to Sb affected chitin- and DNA replication-related processes. Multiomics approaches helped to elucidate the molecular mechanisms of toxicity caused by Sb in soil organisms. Additionally, gene expression profiles exist for well-known model earthworm species such as *Eisenia fetida* (Zwarycz et al., 2016), *Eisenia andrei* (Shao et al. 2020), and *Enchytraeus crypticus* (Amorim et al. 2021). A study by Chai et al. (2020) found that cadmium (Cd) exposure affected enzyme activity, metabolism, oxidative stress, regeneration, and apoptosis pathways in *E. fetida* at the transcriptional level. Their study identified key genes, such as Metallothioneins (MT), Phytochelatins (PCS), Endo-1,4-β-glucanase (EfEG), Superoxide dismutase (SOD), SRY (sex determining region Y)-box 2 (Sox2), SRY-box transcription factor 4b (Sox4b), Apoptosis-stimulating of p53 protein 2-like (ASPP2), and TP53-regulated inhibitor of apoptosis 1 (TRIAP1), which were differentially expressed in response to Cd exposure. The upregulation of MT, PCS, SOD, Sox2, Sox4b, and TRIAP1 suggests a cellular defense mechanism against Cd-induced stress. These findings enhance our understanding of the molecular responses of *E. fetida* to Cd exposure and their potential utility as biomarkers for environmental risk assessment. Genomic and transcriptomic analyses have further contributed to the understanding of the regenerative capacity exhibited by *E. fetida* as demonstrated in the work by Bhambri et al. (2017). They identified key genes involved in regeneration, such as Brachyury and SOX4, which play crucial roles in specifying posterior structures and epithelial-mesenchymal transition (EMT), respectively. The research also revealed the presence of novel genes specific to annelids that contribute to regeneration. Notably, identifying a nerve growth factor (NGF)-like gene in *E. fetida* suggests potential applications for neural regeneration research (Bhambri et al., 2017).

However, such toxicogenomic profiles remain extremely limited for other species that contribute to soil processes, such as carabid beetles, isopods, and soil-dwelling mites (Spurgeon et al., 2008; Brückner et al. 2020). Of soil-dwelling mites, oribatid mites play an integral role in soil health, representing a heterogeneous collection of individuals contributing to soil formation, maintenance, and soil nutrient cycling (e.g., organic matter decomposition) (Wickings and Grandy 2011). Given their abundance, diversity, varied feeding guilds, and limited dispersibility, oribatid mite species are often used as bioindicators of soil contamination (e.g., organic compounds, pesticides, heavy metals pollution) (Behan-Pelletier, 1999, 2003; Gergócs and Hufnagel, 2009; Owojori et al. 2019). Recently, Brückner et al. (2020) published a genome assembly for the soil mite, *Archegozetes longisetosus* Aoki; this species is a model oribatid mite species, representative of tropical habitats, but has not been standardized as a soil test species in so far. The research provides comprehensive genomic and transcriptional analyses of evolutionary and biosynthesis characteristics, providing new insight into soil mites. Similarly, the annotated genome is also available for *Tetranychus urticae*, a polyphageous herbivore mite, often described as an agricultural pest (Grbić et al. 2011). The research by Grbić et al. (2011) provides genetic insight into detoxification mechanisms responsible for the observed pesticide resistance in this organism. The pesticide resistance observed in *T. urticae*, a polyphagous herbivore, can be attributed to the expansion of unique gene families involved in digestion, detoxification, and xenobiotic transport. These expansions include an abundance of cysteine peptidase genes, cytochrome P450 genes, carboxyl/cholinesterases genes, and glutathione S-transferases genes, contributing to their exceptional resilience. Furthermore, these genes exhibit significant changes in expression when spider mites feed on different host plants, with many differentially regulated genes lacking known functions, highlighting the complexity of their detoxification mechanisms (Grbić et al. 2011).

*Oppia nitens* (C.L. Koch, 1836) is an oribatid mite that belongs to the Oppiidae family. This family is one of the largest and most diverse families, with over a thousand known species worldwide (Ohkubo, 2001). *O. nitens* is euryoecious and typically inhabits humus-rich forest and agricultural soils, wetlands, and dry grasslands (Fajana et al., 2019). This mite has been developed as a standard test species for assessing soil contaminants in temperate regions due to its ease of culture, reproduction, varied sensitivity, and broad applicability to diverse soil types (Princz et al., 2010; Owojori and Siciliano, 2012; ISO, 2020). Despite its increased use in soil pollution assessment (Fajana et al., 2019), the lack of genomic information for *O. nitens* limits our understanding of its mechanistic response to pollutants (Princz et al., 2018; Jegede et al., 2019a), its resilience to contaminants (Fajana et al., 2023), and adaptive capacity in different soil habitats (Jegede et al., 2019b). *O. nitens* is sensitive to heavy metals with the ability for the metals to cause transgenerational and multigenerational effects (Owojori and Siciliano, 2012; Jegede et al., 2019a; Fajana et al., 2020). *O. nitens* accumulates certain metals like cadmium at high concentrations (Keshavarz-Jamshidian et al., 2017; Fajana et al., 2019). However, the exact mechanism of Cd bioaccumulation is unknown, but it may be related to the ability of the mite to sequester or compartmentalize metals in specialized proteins (Ludwig et al., 1992; Fajana et al., 2019), therefore, altering metal toxicodynamic in the mites. Fajana et al. (2019) suggested possible molecular pathways like the metallothionein pathways, including phytochelatin-dependent pathway, as the possible mechanism for metal tolerance in *O. nitens*. However, this can only be unraveled if adequate genomic information of the mites is known, which necessitates the complete draft genome of the mite. A detailed knowledge of the mite genome will also aid in understanding gene expression pathways associated with molecular and physiological functions, enabling adaptation to environmental change within their microhabitat, and how the quality of their habitat influences their responses to chemical stressors (Jegede et al., 2019b; Fajana et al., 2021).

Herein, to reduce this significant gap, we present a *de novo* genome assembly of *O. nitens* using a hybrid *de novo* assembly strategy through a combination of Illumina short reads next-generation sequencing and Oxford Nanopore Technologies long reads third-generation sequencing. We annotated the essential genomic elements and protein-coding genes. The contribution of this new genomic data allows for a more detailed assessment of molecular mechanisms associated with toxic responses, detoxification, and metal tolerance, as well as a robust risk assessment of contaminants in soil.

## 2. Materials and Methods

### 2.1 Sample collection and sequencing

#### *2.1.1* DNA extraction

Mite DNA was extracted using the Qiagen DNeasy® blood and tissue kit following the manufacturer’s protocol. Briefly, mites were initially ground using a micro pestle in a 1.5 mL centrifuge tube. Before grinding, the tube was placed in liquid nitrogen for approximately 10 seconds or until bubbling stopped to ensure efficient grinding. The grinding was performed at a low intensity to avoid damaging the DNA.

Due to the mite’s small size, different numbers of well-sclerotized adult mites ranging from 10 to 400 individuals were carefully sorted to optimize extraction protocol. The mites were ground in a series of 10, 20, 50, 100, 200, and 400 mites. It was observed that even 10 mites extracted quantifiable DNA, however, based on yields, it was decided that extracting DNA from 100 mites would be ideal. After grinding the mites, the Qiagen tissue lysis buffer (ATL) and proteinase K were added to the centrifuge tube, followed by incubation at 56 °C for 30 minutes. The tube was vortexed twice during this incubation period to ensure efficient mixing of the sample. Next, the Qiagen buffer (AL) was added to the sample, and the tube was incubated at 56 °C for approximately 5 minutes. This step is important for the removal of contaminants such as proteins and polysaccharides from the sample. The genomic DNA purity and quantity were determined using a Nanodrop 2000 spectrophotometer (Thermo Fisher Scientific Inc., Wilmington, DE) and a Qubit 2.0 fluorometer (Life Technologies Ltd., Paisley, UK). DNA integrity was analyzed using an Agilent TapeStation 4150 (Agilent Technologies, Santa Clara, CA).

#### *2.1.2* Library construction and sequencing

Four libraries were constructed using the Illumina DNA Prep Kit (Illumina, Inc., San Diego, CA, USA). Three libraries were constructed targeting an average insert length of 300 bp, and one targeting an average insert length of 550 bp. The libraries were pooled and 150 bp paired-end reads were generated on an Illumina NextSeq 550 instrument at the University of Saskatchewan Next-Generation Sequencing Facility.

Three additional libraries were constructed for Oxford Nanopore sequencing using the Ligation Sequencing Kit SQK-LSK109 (Oxford Nanopore Technologies, Oxford, UK) and Native Barcoding Kit SQK-NBD114 (Oxford Nanopore Technologies). Nanopore reads were generated on a GridION instrument using an R9.4.1 flow cell at Prairie Diagnostic Services, Inc. (Saskatoon, SK, Canada).

### 2.2 De novo assembly and annotation of O. nitens genome

#### *2.2.1* Genome assembly

A hybrid assembly approach was used for this study. Contigs were assembled from the nanopore reads using Nextdenovo v2.5.0 (https://github.com/Nextomics/NextDenovo) with default settings. Residual adapter and low-quality sequences were trimmed from the Illumina short reads using fastp v0.20.0 with default settings (Chen et al., 2018). The trimmed reads were aligned to the nextdenovo contigs using bwa-mem2 v2.2.1 (https://github.com/bwa-mem2/bwa-mem2) with default settings, and sequencing errors corrected using the aligned reads and Pilon v1.23 (Walker et al., 2014). Additional scaffolding, error correction, and gap-filling was performed using longstitch v1.0.1 (Coombe et al., 2021) with the consensus-polished reads output by nextdenovo and default settings aside from: k=32, w=200.

#### *2.2.2* Genome annotation

The genome sequence records for *O. nitens* RefSeq assembly (GCF_028296485.1) were annotated by the National Center for Biotechnology Information (NCBI) Eukaryotic Genome Annotation Pipeline (EGAP), an automated pipeline that annotates genes, transcripts, and proteins on draft and finished genome assemblies (https://www.ncbi.nlm.nih.gov/refseq/annotation_euk/process/). Transcript information from RNA sequencing (RNA-Seq) data (described below) was provided to assist in gene prediction.

Blast2GO PRO v3.1.9 (Blast2GO - Götz et al., 2008) was used to obtain gene functional classification based on Gene Ontology (GO) terms in the “biological process,” “molecular function,” and “cellular component” categories. Kyoto Encyclopedia of Genes and Genomes (KEGG) Analysis was also performed for functional annotation. KEGG orthology terms were assigned using GhostKOALA (Kanehisa et al. 2016) (GHOSTX searches for KEGG Orthology And Links Annotation) to obtain KEGG orthology terms. In contrast to traditional BLAST searches, GhostKOALA demonstrates approximately 100-fold greater efficiency in identifying remote homologs, achieved through the utilization of suffix arrays (Suzuki et al., 2014).

#### *2.2.3* De novo assembly and annotation of *O. nitens* mitochondrial genome

The mitochondrial genome was assembled and annotated with MitoZ v3.6 (Meng et al., 2019) from the DNA short reads (all read files concatenated together) used to generate the *O. nitens* draft genome using MEGAHIT for assembly and the Arthropoda clade, and kmers 59 79 99 119 141.

### 2.3 RNA-seq data generation

#### *2.3.1* Soil exposure for preliminary transcriptomics test

Cadmium oxide (CdO) (≥ 99.99% trace metal basis) (Sigma-Aldrich, Canada) was used as the test chemical. Four test soils were used for the exposure assays: one high habitat quality natural soil; one low habitat quality natural soil; OECD artificial standard soil; and LUFA 2.2 (LUFA Speyer, Germany) natural standard soil. The high- and low-quality natural soils were collected from western Canada and grouped into quality categories based on the performance of standard test organisms as described in Jegede et al. (2019b).

Age-synchronized adult mites were exposed to Cd-spiked soil for 16 days at the following nominal concentrations: 0, 215, and 645 mg/kg of dry soil. The concentrations were based on the results from our previous reproduction test to derive sublethal concentrations (Fajana et al. 2019). Test vessels consisted of control and Cd-spiked soils with six replicates per treatment; each replicate had 15 mites. After exposure, mites were collected by heat extraction (modified Berlese-Tullgren heat extractor set at 32 ℃) and rinsed, transferred to 1.5 mL centrifuge tubes, snap frozen with liquid nitrogen, and stored at -80 ◦C until RNA extraction. Samples from the highest concentration (645 mg/kg) were excluded due to mortality, therefore, only the control and 215 mg/kg dry soil treatments were used for RNA extractions for each soil type.

#### *2.3.2* Total RNA extraction and cDNA library construction

Prior to extraction, liquid nitrogen was added to the collection tubes, and the mites were ground into a powder using micro pestles. Total RNA was isolated from each sample using the RNeasy® mini kit (Qiagen) and RNase-free DNase set (Qiagen), following the manufacturer’s protocol. RNA concentration, quality, and integrity were checked using Qubit 2.0, Nanodrop, and Agilent 2100, respectively.

Strand-specific RNA-Seq libraries were constructed from 16 total RNA samples using the NEBNext Ultra II Directional RNA Library Prep Kit for Illumina (New England Biolabs, Ipswich, MA, USA) with NEBNext Poly(A) mRNA Magnetic Isolation Module (New England Biolabs). To gain insight into the nonpolyadenylated fraction of the transcriptome, one additional library per sample was constructed using the Zymo-Seq RiboFree Total RNA Library Kit (Zymo Research, Irvine, CA, USA). Each library type was pooled and 150 bp paired-end reads generated on an Illumina NextSeq 550 instrument at the University of Saskatchewan Next-Generation Sequencing Facility. RNA-Seq data was used to provide more gene coverage in gene prediction.

### 2.4 Ortholog Clustering and Gene Family Analysis

We inferred orthogroups of 7 mite species using OrthoFinder v2.4.0 (Emms and Kelly, 2019). The annotated protein set from *O. nitens* (BioProject accession: PRJNA905106) and protein sequences from *Dermatophagoides farinae* (PRJNA512594), *Galendromus* (*Metaseiulus*) *occidentalis* (PRJNA62309), *Medioppia subpectinata* (PRJEB39968), *Oppiella nova* (PRJEB39968), *Tetranychus urticae* (PRJEA71041), and *Varroa destructor* (PRJDB6279). The protein sequences were downloaded from the NCBI database. Gene family evolution (expansions and contractions) was estimated using CAFÉ v5.0. (Mendes et al., 2020) lambda parameter to calculate birth and death rates.

## 3. Results and discussion

### 3.1 De novo genome assembly

Our final draft assembly of *O. nitens* genome consists of 65 scaffolds/contigs of 125.4Mb, a scaffold/contig N50 length of 4.4Mb which demonstrates contiguity, and 24.5% GC content (Table 1). Data obtained from genome completeness analysis by BUSCO assessment against arthropod set (n=2934) showed that 93.5% of the predicted genes on the assembled genomic sequences were complete and single copy, 3.4% were complete and duplicated, and only a few were fragmented (0.5%) or missing (2.6%) BUSCO orthologs. This indicates the high completeness and low redundancy of our assembly. The assembled genome size of *O. nitens* is smaller than that of other acariform and mesostigmatid mites, ticks, and spiders (Bast et al. 2016; Dong et al. 2017; Dong et al. 2018; Grbić et al. 2011; Gulia-Nuss et al. 2016; Hoy et al. 2016; Schwager et al. 2017). Among other acariform mites, *O. nitens’s* genome (Table 1) falls on the small end and shares this feature with other arthropod species with smaller genomes, such as the spider mite, *T. urticae* (90Mb) (Grbić et al. 2011).

**Table 1.** Properties of *O. nitens* genome

| **De novo assembly** |  |
| --- | --- |
| Genome size (Mb) | 126.1 |
| Total ungapped length (Mb) | 125.4 |
| No. scaffolds | 65 |
| N50 length (Mb) | 4.4 |
| Scaffold L50 | 7 |
| GC (%) | 24.5 |
| Complete | 93.5 |
| Complete and duplicated | 3.4 |
| Fragmented | 0.5 |
| Missing | 2.6 |
| Coverage | 440.0x |
| **Genome structure** |  |
| Total number of genes | 15,291 |
| Average gene length (bp) | 3,708 |
| Number of protein-coding genes | 14,938 |
| Non-coding | 353 |
| Exons | 72,268 |
| Average exon length | 355 |
| Introns | 56,531 |
| Average intron length | 616 |

### 3.2 Mitochondrial genome

Our assembled mitochondrial (mt) genome of *O. nitens* is 13,867 base pairs (bp) long. It has 13 protein-coding genes, 12 transfer RNA (tRNA) genes, and 2 rRNA genes, totaling 27 genes (Fig. 1). These include genes encoding for NADH dehydrogenase subunits, cytochrome b, cytochrome c oxidase subunits, ATP synthase subunits, and various tRNAs and rRNAs. The mitogenome of Sarcoptiformes contains a conserved set of 37 genes which includes 13 protein-coding genes, 2 rRNAs genes and 22 tRNA genes (Salinas-Giegé et al., 2015). In *O. nitens*, however, part of the NADH dehydrogenase 4L (ND4L) gene, and 10 tRNA genes: Alanine, Arginine, Cysteine, Histidine, Methionine, Phenylalanine, two Serine tRNAs, Tyrosine, and Valine were not identified.

**Figure 1.**
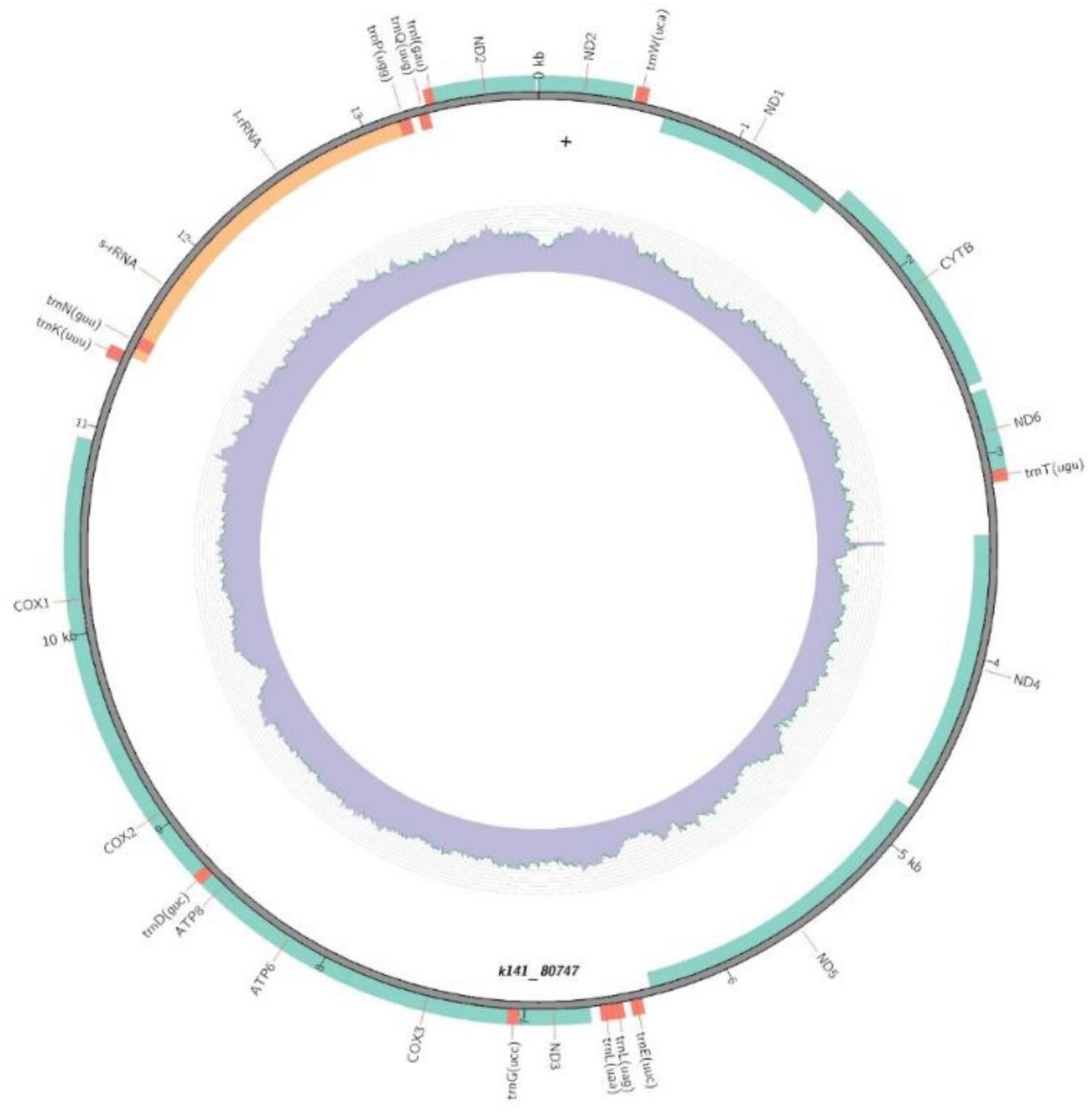
Circular representation of the *O. nitens* mitochondrial genome generated via MitoZ. The outer ring displays the annotated genes, including protein-coding genes (colored teal), tRNA genes (colored red), and rRNA genes (colored orange). The inner ring illustrates the GC content across the genome.

Many mitogenomes, including those of oribatid mites, exhibit the absence of one or more tRNA genes, with some species lacking all tRNAs entirely (Schneider, 2011; Salinas-Giegé et al., 2015; Schäffer et al., 2018). Four species within Acariformes demonstrate a marked reduction in mt-tRNA genes: *Steganacarus magnus* (16 tRNAs) (Domes et al., 2008), *Sarcoptes scabiei* (three tRNAs) (Mofiz et al., 2016), and two *Tyrophagus* species (two tRNAs) (Yang and Li, 2016). The observed absence of 10 mt-tRNAs in *O. nitens* may reflect such evolutionary processes, where the functional role of these genes could be compensated by nuclear-encoded cytoplasmic tRNAs that perform similar roles within the mitochondria (Clayton, 1992; Adams and Palmer, 2003).

While gene loss, particularly tRNA gene loss, is not uncommon in mitochondrial genomes, the absence of genes in *O. nitens’* mitogenome may be partly due to limitations in current predictive tools rather than their actual loss. Accurately identifying mt-tRNA genes is inherently challenging, especially when they exhibit unconventional secondary structures, deviate from standard codon-anticodon rules, or undergo post-transcriptional modifications (Huot et al., 2014). Despite the availability of advanced bioinformatics tools, such complexities can hinder reliable prediction, leading to potential under-detection or misidentification of mt-tRNAs (Huot et al., 2014). Structural deviations from canonical forms may render these mt-tRNAs difficult to detect through traditional genome annotation approaches, especially in mite species such as *T. urticae* where mitochondrial tRNA genes are known to be highly degenerate (Warren and Sloan, 2021).

Studies such as Fang et al. (2020) argue against the loss of mt-tRNA genes after identifying tRNA genes in *Tyrophagus putrescentiae* that were initially reported as missing, demonstrating the potential for predictive tools to overlook atypical tRNA structures. Similarly, Xue et al. (2018) questioned earlier findings of mt-tRNA gene loss in Sarcoptiform mites, suggesting that such genes might remain undetected due to the inherent challenges in identifying highly modified or unusual tRNAs. These findings emphasize the need for careful annotation and raise the possibility that the mt-tRNAs in *O. nitens* may still be present but undetected due to the complexity of their structure. A combined multi-software and manual annotation approach, as applied by Zhan et al. (2021), could provide greater accuracy and help determine whether the missing genes are indeed absent or simply undetected due to limitations in current annotation methods.

### 3.3 Gene Prediction and functional annotation of predicted genes

We used the NCBI EGAP to predict complete genes in the *O. nitens* genome. The analysis resulted in 15,291 total genes, of which 14,938 were protein-coding. The mean lengths of genes, exons, and introns were 3,708, 355, and 616 bp, respectively. We identified 393 non-coding (ncRNAs): 52 rRNAs, 21 small nuclear RNAs (snRNAs), 27 miscRNAs, 161 long ncRNAs (lncRNAs), 18 small nucleolar RNA (snoRNA), and 114 tRNAs (Supplementary Table S1, Supplementary Material).

InterproScan and eggNOG functional annotations on Blast2GO assigned protein domains to 14,789 genes, 13,246 gene ontology (GO) terms (Fig 2.), and 8,753 Reactome pathways. Fourteen biological processes were identified at level 3 GO with the most abundant being: “organic substance metabolic process” (15%), “primary metabolic process” (15%), “nitrogen compound metabolic process” (14%), and “cellular metabolic process” (13%). Among the GO terms identified for molecular functions, the majority were constituted by: “organic cyclic compound binding” (22%), “small molecule binding” (17%), “protein binding” (16%), and “hydrolase activity” (14%). For cellular component, a total of seven GO terms were identified with the most represented being: “intracellular anatomical structure” (29%), “organelle” (24%), “cytoplasm” (19%), and “membrane” (13%) (Fig. 2).

**Figure 2.**
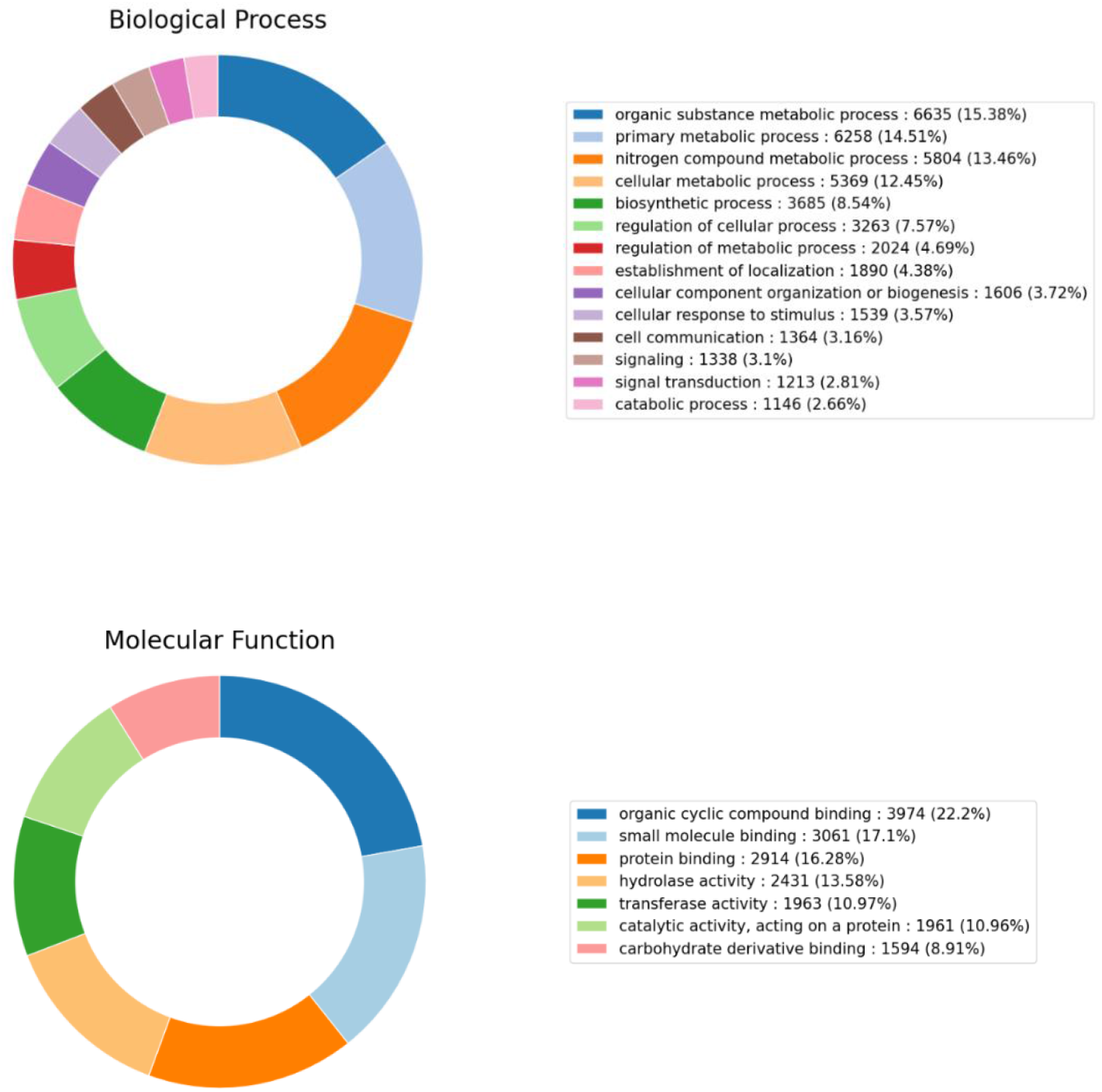
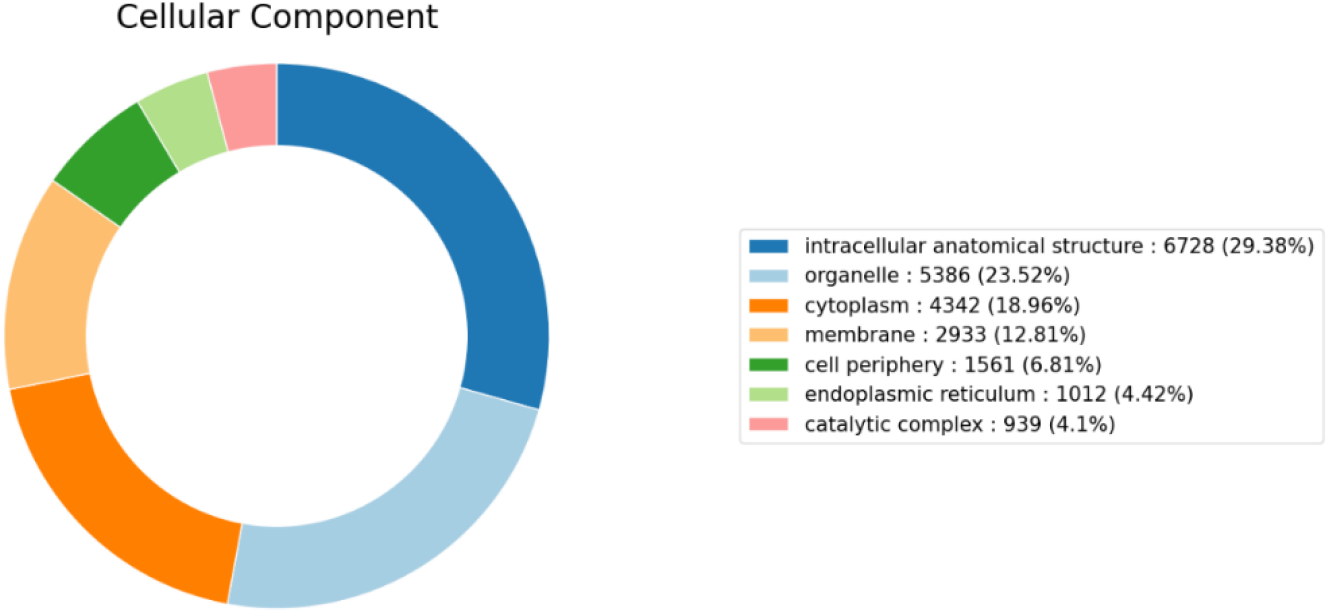
Gene ontology classification of the genes predicted from *O. nitens* genome assembly: A gene ontology (GO) term was assigned to each gene, which was summarized into three main GO categories (A: biological process, B: cellular component, and C: molecular function).

In addition, we ran a GhostKOALA (KEGG Orthology And Links Annotation) analysis to characterize the genes functionally (Kanehisa et al. 2016). Over 60.5% of coding sequences (CDS) entries were assigned functional characteristics. Most genes were either metabolic genes or genes related to genetic information processing, while the remaining genes were assigned to other functional categories (Fig. 3).

**Figure 3.**
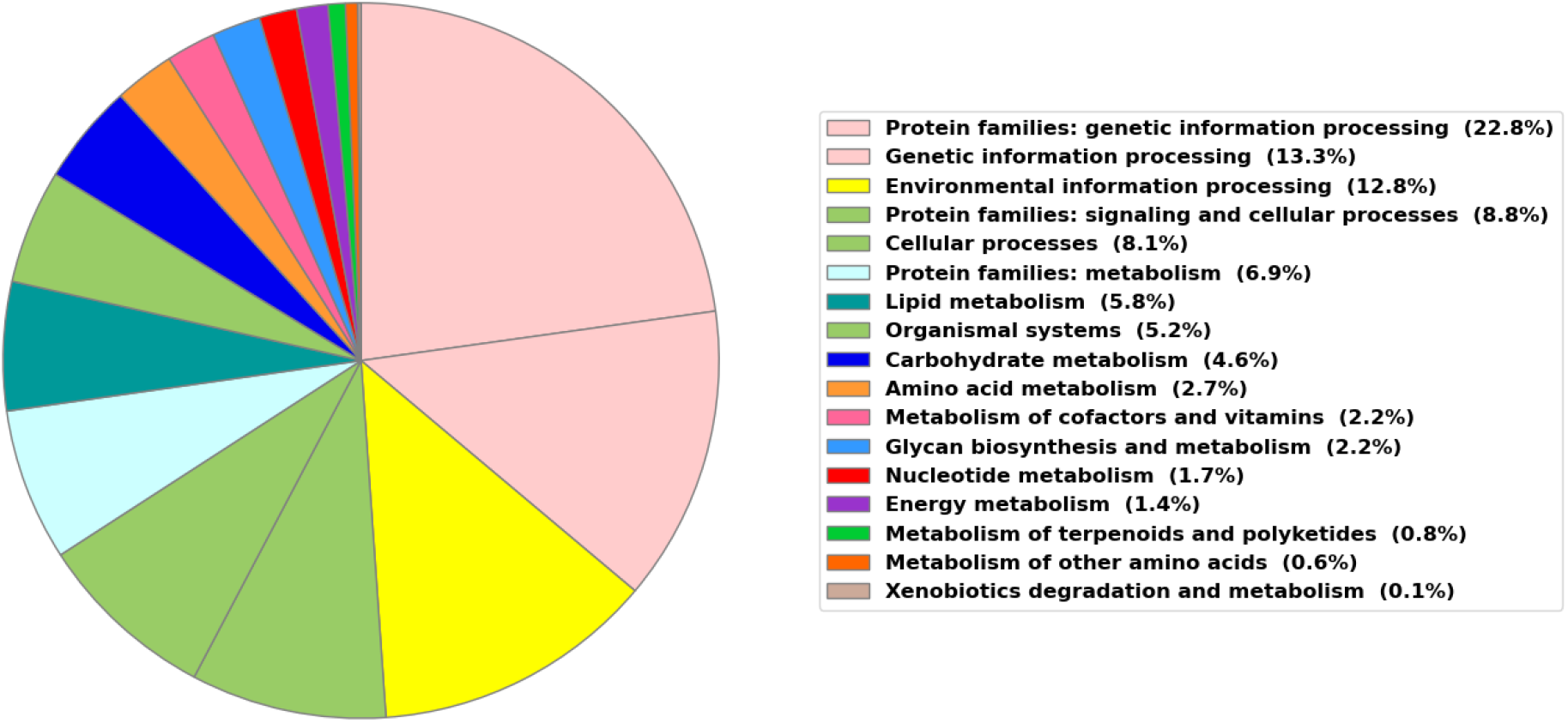
Kyoto encyclopedia of genes and genomes (KEGG) functional categories and pathways for predicted proteins of *O. nitens* using GhostKOALA.

The annotation of *O. nitens*’ genome reveals its extensive metabolic capabilities, with a significant portion of genes involved in organic substance metabolic processes, biosynthetic activities, and cellular localization (Fig. 2). These processes are fundamental to the organism’s ability to metabolize nutrients and maintain cellular organization, which are essential for its survival in its widely distributed natural habitats (Fajana et al., 2019). The biosynthetic processes and the substantial representation of genes involved in cell communication and catabolic activities highlight *O. nitens*’ capacity to synthesize critical biomolecules and recycle nutrients efficiently. KEGG functional categorization further reveals the metabolic and regulatory pathways vital for environmental adaptation. The prominence of genes related to genetic information processing (22.8%, 13.3%) and environmental information processing pathways (12.8%) in the KEGG analysis reflects *O. nitens*’ robust mechanisms for genomic maintenance (DNA replication and repair) and environmental sensing. These pathways (Table 2) are vital for *O. nitens* as it navigates and responds to environmental stressors, such as metal contamination, which can impose significant oxidative stress (Owojori and Siciliano, 2012). For instance, upon detecting the presence of heavy metals, signal transduction pathways such as Mitogen-Activated Protein Kinase (MAPK), Phosphoinositide 3-Kinase (PI3K)-Akt, and Nuclear factor-erythroid 2 related factor 2 (Nrf2) are activated, leading to the upregulation of genes involved in detoxification, and stress response (Kumagai and Sumi, 2007). Membrane transporters, e.g., ATP-binding Cassette (ABC) Transporters, can facilitate the efflux of metal ions or metal-chelate complexes from the cell preventing toxicity (Jalmi, 2022). Genes related to other pathways involved in stress response and toxicity adaptation, such as Carbohydrate metabolism (4.6%) – associated with habitat quality, Metabolism of other amino acids (0.6%) – associated with glutathione metabolism and important for phytochelatin synthesis, and Xenobiotics degradation and metabolism (0.1%) are also represented (Fig 3., Table 2).

**Table 2.** Relevant KEGG functional categories and associated pathways related to stress response and toxicity adaptation

| KEGG functional categories | Associated pathways |
| --- | --- |
| Genetic Information Processing | Replication and repair |
|  | Base excision repair<br>Nucleotide excision repair<br>Mismatch repair<br>Homologous recombination |
| Environmental Information Processing | **Membrane transport**<br>ABC transporters<br>**Signal transduction**<br>MAPK signaling pathway<br>PI3K-Akt signaling pathway<br>mTOR signaling pathway<br>TGF-beta signaling pathway<br>NF-kappa B signaling pathway<br>HIF-1 signaling pathway<br>JAK-STAT signaling pathway<br>Wnt signaling pathway |
| Lipid metabolism | Arachidonic acid metabolism<br>Steroid hormone biosynthesis<br>Sphingolipid metabolism<br>Fatty acid metabolism |
| Carbohydrate Metabolism | Glycolysis / Gluconeogenesis<br>Citrate cycle (TCA cycle)<br>Pentose phosphate pathway |
| Energy Metabolism | Oxidative phosphorylation |
| Metabolism of Cofactors and Vitamins | Ubiquinone and other terpenoid-quinone biosynthesis |
| Metabolism of Other Amino Acids | Glutathione metabolism |
| Cellular Processes | Autophagy<br>Apoptosis |
|  | Endocytosis |
|  | Lysosome |
| Organismal Systems | Immune system pathways<br>Cytokine-cytokine receptor interaction<br>Complement and coagulation cascades<br>Inflammatory mediator regulation of TRP channels |
| Xenobiotics Degradation and Metabolism | Benzoate degradation<br>Aminobenzoate degradation<br>Fluorobenzoate degradation<br>Chloroalkane and chloroalkene degradation<br>Chlorocyclohexane and chlorobenzene degradation<br>Toluene degradation<br>Xylene degradation<br>Nitrotoluene degradation<br>Ethylbenzene degradation<br>Styrene degradation<br>Atrazine degradation<br>Caprolactam degradation<br>Bisphenol degradation<br>Dioxin degradation<br>Naphthalene degradation<br>Polycyclic aromatic hydrocarbon degradation<br>Furfural degradation<br>Steroid degradation<br>Drug metabolism - cytochrome P450<br>Metabolism of xenobiotics by cytochrome P450 |

Note: **PI3K-Akt**: Phosphoinositide 3-Kinase-Akt signaling pathway, **mTOR**: Mammalian Target of Rapamycin signaling pathway, **TGF-beta**: Transforming Growth Factor-beta signaling pathway, **NF-kappa B**: Nuclear Factor-kappa B signaling pathway, **HIF-1**: Hypoxia-Inducible Factor 1 signaling pathway, **AK-STAT**: Janus Kinase-Signal Transducer and Activator of Transcription signaling pathway, **Wnt**: Wingless-Integration signaling pathway

### 3.4 Genome size and GC content

Mites typically have small genomes (Gregory and Young, 2020), but it is interesting to observe genome size variation even among smaller genomes. With a 126-Mbp genome, *O. nitens* has a streamlined genome compared to close Oribatida relatives; smaller than *Oppiella nova* (197 Mbp) and *Oppiella subspectinata* (210 Mbp). It is also smaller than other mite species, such as *Stratiolaelaps scimitus* (427 Mbp), but larger than *T. urticae* (90 Mbp), which is one of the smallest arthropod genomes known so far (Grbic et al., 2011) (Fig. 4). Genome streamlining is an adaptive and evolutionary mechanism that allows organisms to dispense unnecessary genes from their genome to adapt to extreme environmental conditions (Sengupta et al., 2024).

**Figure 4.**
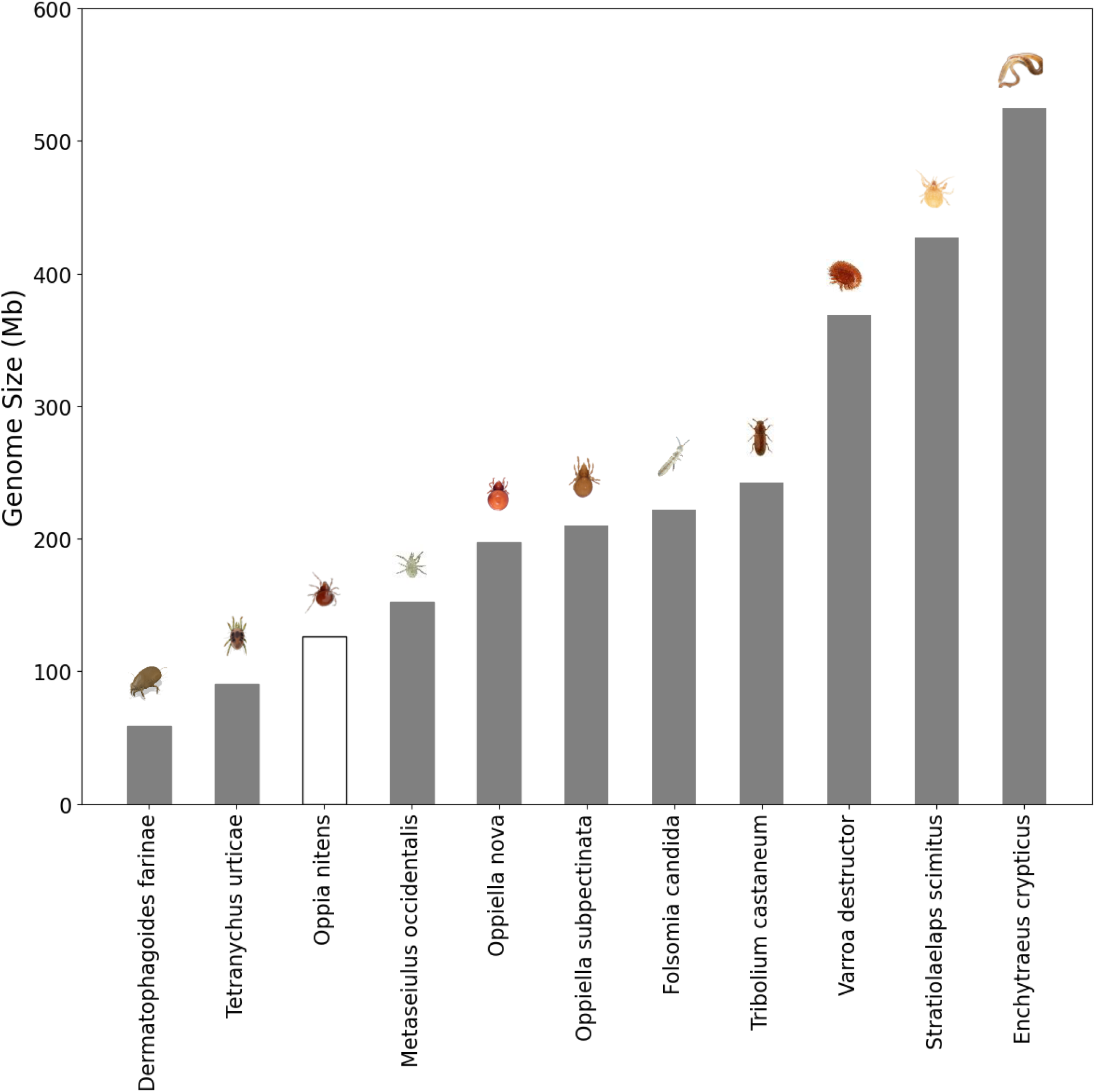
Genome size comparison between nine mite species, *F. candida*, and *E. crypticus*. The bar chart shows the genome sizes of different species, highlighting *O. nitens* in black with a streamlined genome size of 126 Mb (Accession information in supplementary Table S2).

Genome size variation occurs within and among different taxonomic levels of plants and animals, giving rise to several hypotheses for the factors driving the evolution of both small and large genomes (Gregory, 2005; Lynch and Walsh, 2007). In eukaryotes, larger body size was hypothesized to correlate with larger genomes (Gregory, 2002). For example, in rotifers, the genome size correlated positively with body size (Stelzer et al., 2021). However, comparing two Oribatid mite species, this was not the case as *O. nitens* has a larger body size (length: 510 μm) (Fajana et al. 2019) than *O. nova* (length: 231 and 297μm) (von Saltzwedel et al. 2014) but has a relatively smaller genome and lower number of genes compared to *O. nova* (Fig. 5). *O. nova* is a free-living oribatid mite that mainly feeds on fungal hyphae and inhabits the upper organic-rich soil layer subject to environmental variation. It is parthenogenetic but also iteroparous and has a 23-day life cycle (von Saltzwedel et al. 2014). Just like *O. nitens, O. nova* possesses life history attributes generally present in oribatid mites, such as low metabolic rates, which enables them to survive when food resources are limited (Norton 1994). With organisms possessing smaller genomes being more nutrient efficient (Giovannoni et al., 2014), having a small genome might be a useful evolutionary adaptation strategy for oribatid mites to adapt to their environment. In the case of *O. nitens* with a more streamlined genome, they might be even more adaptable to challenging conditions e.g., poor habitat quality conditions or chemical stress.

**Figure 5.**
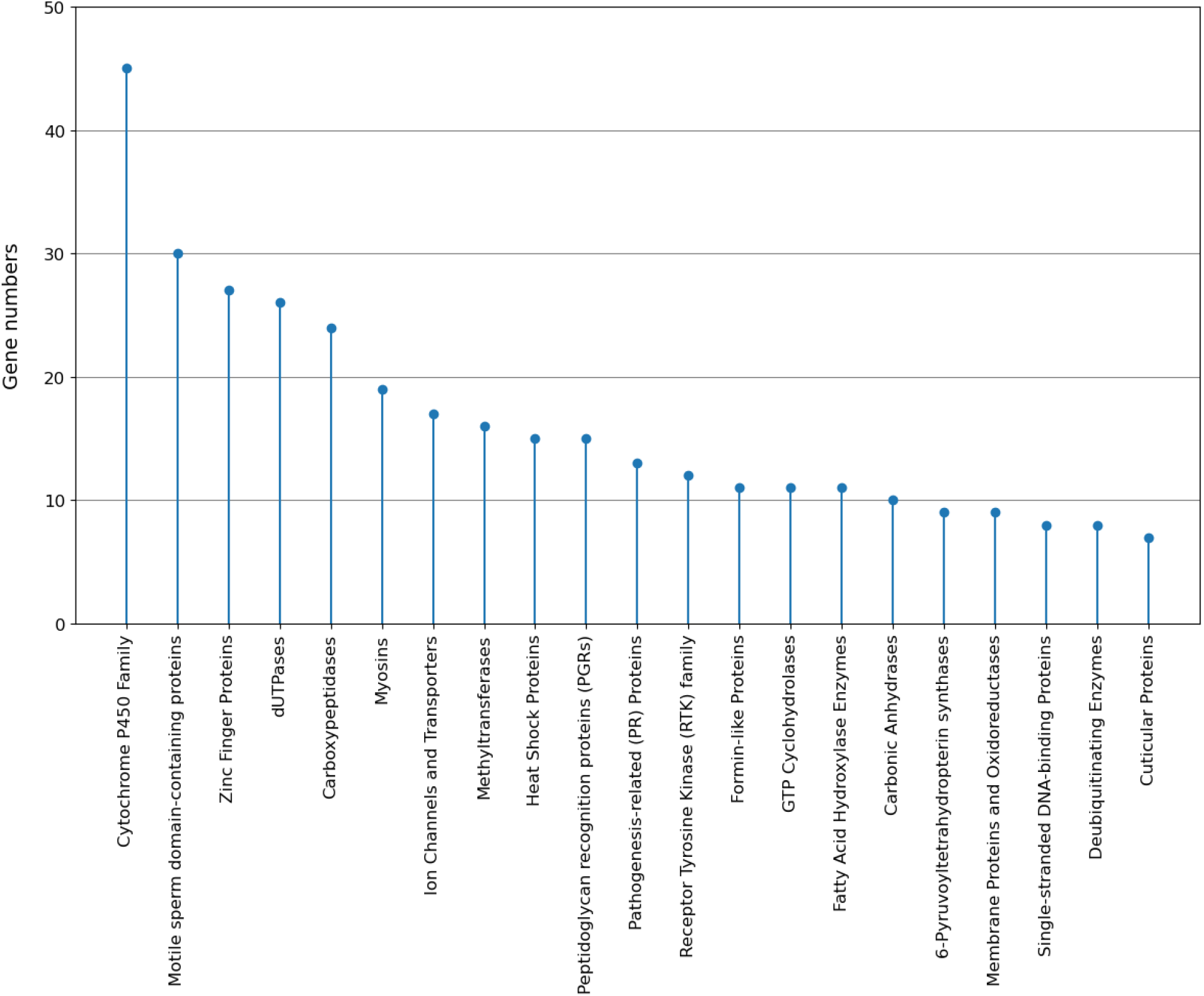
Top significantly expanded *O. nitens* gene families including their gene numbers.

The genomic scaffold’s nucleotide composition indicates that the draft sequence consists of GC at 24.5% of total bases in *O. nitens* genome, a value much below the range (30% – 45%) observed not just among other oribatid mites, but also in other mite species and test soil species (Table 3). The notably low GC content in *O. nitens* suggests a genome that may be more flexible but potentially less stable than its relatives and other species (Table 3). GC-rich regions are known for their stronger bonding between guanine and cytosine bases, which provides greater stability to the DNA structure (Smarda et al., 2014). This increased stability can influence the efficiency and fidelity of DNA replication and repair processes (Smarda et al., 2014). In the case of *O. nitens*, the lower GC content might indicate a genome more prone to certain types of mutations, such as AT-biased mutations (Wu et al., 2012), which could be a factor for adaptation to rapidly changing or stressful environments. *O. nitens’* low GC content is surprisingly close to many *Plasmodium* species (17% – 23%), noted as being one of the most AT-rich eukaryotic organisms (Videvall, 2018). Their low GC content has been linked to their unique host-pathogen interactions and life cycle adaptations (Hamilton et al., 2017). While *O. nitens* is not parasitic, its low GC content may reflect similar evolutionary pressures, such as the need to rapidly adapt to challenging environments.

**Table 3.** GC content variation between close *O. nitens* relatives, two other mite species, *F. candida* (Collembolan) and *E. crypticus* (Enchytraeid)

| Species |  | GC content (%) |
| --- | --- | --- |
| *Oppia nitens* | Oribatid mite | 24.5 |
| *Oppiella nova* | Oribatid mite | 30.2 |
| *Oppiella subpectinata* | Oribatid mite | 30.8 |
| *Stratiolaelaps scimitus* | Mesostigmatid mite | 45.9 |
| *Tetranychus urticae* | Tetranychid mite | 32.5 |
| *Folsomia candida* | Collembolan | 37.5 |
| *Enchytraeus crypticus* | Enchytraeid | 35.4 |

The evolutionary implications of this lower GC content are particularly intriguing. It suggests that *O. nitens* may have evolved under selective pressures that differ from those experienced by its relatives with higher GC content. The need for rapid replication and potentially lower energetic costs of DNA synthesis in environments where resources are limited or where rapid population growth is advantageous could have driven the evolution of a genome with a lower GC content in *O. nitens* (Wu et al., 2012).

### 3.5 Gene family expansions

Gene families were identified among 7 mite species with OrthoFinder. A total of 92.2% (123,985) genes were clustered into 17,487 orthogroups (gene families). Among them, 3,923 families were shared by all 7 species, and 294 were single-copy ones; 4913 families and 19,358 genes were species-specific. In *O. nitens*, 16,969 (93.2%) genes were clustered into 15,814 gene families, and 495 families and 2255 genes were species-specific (Supplementary Table S3). We analyzed gene family evolution (gain and loss) using CAFÉ, and the estimated gene birth rate (lambda) was 0.586964. A total of 396 gene families experienced significant expansion or contraction events across the tree with a family-wide P-value <0.05 (Supplementary Table S4). The top largest expanded families are shown in Figure 5. Some of these include Cytochrome P450, Zinc finger proteins, Heat shock proteins, Pathogenesis-related proteins (Golgi-associated plant pathogenesis-related proteins), Deubiquitinating enzymes, and Cuticular proteins.

The most expanded family, cytochrome P450s, plays an important role in the metabolism of xenobiotics and endogenous compounds (Feyereisen, 2012). Zinc finger proteins are transcription factors that bind to DNA and regulate gene expression. They are involved in numerous cellular processes, including DNA repair, apoptosis, and the response to oxidative stress. These proteins are critical in upregulating genes involved in detoxification and stress response. For example, they may activate the transcription of metallothioneins and antioxidant enzymes, essential for binding and neutralizing metal ions, thus protecting the organism from metal-induced oxidative damage (Laity et al., 2001). Heat shock proteins are molecular chaperones rapidly deployed in stress response to assist in protein folding, prevent protein aggregation, and help in the degradation of damaged proteins (Li and Srivastava, 2003). Pathogenesis-related proteins (Golgi-associated plant pathogenesis-related proteins) contribute to pathogen defense, help maintain protein homeostasis, and modulate immune responses (Jain and Khurana, 2018). Deubiquitinating enzymes are essential in maintaining protein homeostasis and responding to cellular stress (Amerik and Hochstrasser, 2004). It is interesting to find the expansion of cuticular proteins in the *O. nitens* genome. Previous studies have shown the ability of *O. nitens* to bioaccumulate metals (Cd) in their cuticle (Keshavarz-Jamshidian et al., 2017; Fajana et al., 2019). The expansion of cuticular proteins provides evidence of the ability of *O. nitens* to chelate and bioaccumulate heavy metals in their cuticle, allowing them to tolerate high concentrations of metals in their environment.

The expansion of these families is essential for adaptation to the complicated soil environment. Overall, *O. nitens* exerts some remarkable features concerning genetic adaptation to environmental stress. The resilience of *O. nitens* to chemical stress, especially heavy metals, has been reported in previous studies (Fajana et al., 2023). Gene family evolution provides essential evidence, supporting genetic mechanisms of ecological adaptions for *O. nitens*. Further studies e.g., changes in gene expression after exposure to chemical stress are required to observe the expression patterns of these expanded genes.

## 4. Conclusions

Using a combination of Nanopore Oxford and Illumina reads, we report the first high-quality draft genome for *O. nitens*, resulting in a 126 Mbp assembly with contiguity and completeness, and low GC content of 24.5%. The genome appears relatively streamlined when compared to other annotated genomes of Oribatid mites as well as other chelicerate species available in public databases. This compact genome, lower GC content, and a significant number of expanded gene families related to detoxification and stress response provide genomics-based evidence for *O. nitens’* evolutionary strategy for survival in challenging environments. These findings provide valuable insights into the genetic and molecular mechanisms that enable *O. nitens* to thrive in contaminated soils, making it an important model for studying environmental stress response and soil health.

## Acknowledgments

This research is supported by NSERC (Natural Science and Engineering Research Council of Canada) grant that was awarded to S.D. Siciliano. The authors declare no conflict of interest.

## Data Availability

Raw sequencing data and genome assembly of *O. nitens* have been deposited at the NCBI under BioProject accession PRJNA905106.

## References

1. Adams, K. L., & Palmer, J. D. (2003). Evolution of mitochondrial gene content: Gene loss and transfer to the nucleus. Molecular Phylogenetics and Evolution, 29(3), 380–395.

2. Amerik, A. Y., & Hochstrasser, M. (2004). Mechanism and function of deubiquitinating enzymes. Biochimica et Biophysica Acta (BBA) - Molecular Cell Research, 1695(1-3), 189–207.

3. Amorim, M. J., Gansemans, Y., Gomes, S. I., Van Nieuwerburgh, F., & Scott-Fordsmand, J. J. (2021). Annelid genomes: *Enchytraeus crypticus*, a soil model for the innate (and primed) immune system. Lab Animal, 50(10), 285–294.

4. Bast, J., Schaefer, I., Schwander, T., Maraun, M., Scheu, S., & Kraaijeveld, K. (2016). No accumulation of transposable elements in asexual arthropods. Molecular Biology and Evolution, 33(3), 697–706.

5. Behan-Pelletier, V. M. (1999). Oribatid mite biodiversity in agroecosystems: Role for bioindication. Agriculture, Ecosystems & Environment, 74(1-3), 411–423.

6. Behan-Pelletier, V. M. (2003). Acari and Collembola biodiversity in Canadian agricultural soils. Canadian Journal of Soil Science, 83(Special Issue), 279–288.

7. Bhambri, A., Dhaunta, N., Patel, S. S., Hardikar, M., Srikakulam, N., Shridhar, S., Vellarikkal, S., Suryawanshi, H., Pandey, R., Jayarajan, R., & Verma, A. (2017). Insights into regeneration from the genome, transcriptome, and metagenome analysis of *Eisenia fetida*. BioRxiv, p.180612.

8. Brinke, A., & Buchinger, S. (2017). Toxicogenomics in environmental science. *In vitro Environmental Toxicology - Concepts*, Application and Assessment, 159–186.

9. Brückner, A., Barnett, A. A., Antoshechkin, I. A., & Kitchen, S. A. (2020). Molecular evolutionary trends and biosynthesis pathways in the Oribatida revealed by the genome of *Archegozetes longisetosus*. BioRxiv, 2020-12.

10. Chai, L., Yang, Y., Yang, H., Zhao, Y., & Wang, H. (2020). Transcriptome analysis of genes expressed in the earthworm *Eisenia fetida* in response to cadmium exposure. Chemosphere, 240, 124902.

11. Chen, G., de Boer, T. E., Wagelmans, M., van Gestel, C. A., van Straalen, N. M., & Roelofs, D. (2014). Integrating transcriptomics into triad-based soil-quality assessment. Environmental Toxicology and Chemistry, 33(4), 900–909.

12. Chen, S., Zhou, Y., Chen, Y., & Gu, J. (2018). fastp: An ultra-fast all-in-one FASTQ preprocessor. Bioinformatics, 34(17), i884–i890.

13. Clayton, D. A. (1992). Transcription and replication of animal mitochondrial DNAs. International Review of Cytology, 141, 217–232.

14. Coombe, L., Li, J. X., Lo, T., Wong, J., Nikolic, V., Warren, R. L., & Birol, I. (2021). LongStitch: High-quality genome assembly correction and scaffolding using long reads. BMC Bioinformatics, 22, 1–13.

15. Crain, C. M., & Bertness, M. D. (2006). Ecosystem engineering across environmental gradients: Implications for conservation and management. Bioscience, 56, 211–218.

16. de Boer, M. E., de Boer, T. E., Mariën, J., Timmermans, M. J., Nota, B., van Straalen, N. M., Ellers, J., & Roelofs, D. (2009). Reference genes for QRT-PCR tested under various stress conditions in *Folsomia candida* and *Orchesella cincta* (Insecta, Collembola). BMC Molecular Biology, 10, 1–11.

17. de Boer, T. E., Janssens, T. K., Legler, J., van Straalen, N. M., & Roelofs, D. (2015). Combined transcriptomics analysis for classification of adverse effects as a potential end point in effect-based screening. Environmental Science & Technology, 49(24), 14274–14281.

18. Decaëns, T., Jiménez, J. J., Gioia, C., Measey, G. J., & Lavelle, P. (2006). The values of soil animals for conservation biology. European Journal of Soil Biology, 42, 23–38.

19. Domes, K., Maraun, M., Scheu, S., & Cameron, S. L. (2008). The complete mitochondrial genome of the sexual oribatid mite *Steganacarus magnus*: Genome rearrangements and loss of tRNAs. BMC Genomics, 9, 1–13.

20. Dong, X., Armstrong, S. D., Xia, D., Makepeace, B. L., Darby, A. C., & Kadowaki, T. (2017). Draft genome of the honeybee ectoparasitic mite, *Tropilaelaps mercedesae*, is shaped by the parasitic life history. Gigascience, 6(3), gix008.

21. Dong, X., Chaisiri, K., Xia, D., Armstrong, S. D., Fang, Y., Donnelly, M. J., Kadowaki, T., McGarry, J. W., Darby, A. C., & Makepeace, B. L. (2018). Genomes of trombidid mites reveal novel predicted allergens and laterally transferred genes associated with secondary metabolism. GigaScience, 7(12), giy127.

22. Emms, D. M., & Kelly, S. (2019). OrthoFinder: Phylogenetic orthology inference for comparative genomics. Genome Biology, 20, 238.

23. Fábio, K. M., Vanderpool, D., Fulton, B., & Hahn, M. W. (2020). CAFE 5 models’ variation in evolutionary rates among gene families. *Bioinformatics*, btaa1022.

24. Faddeeva-Vakhrusheva, A., Derks, M. F., Anvar, S. Y., Agamennone, V., Suring, W., Smit, S., … & Roelofs, D. (2016). Gene family evolution reflects adaptation to soil environmental stressors in the genome of the collembolan *Orchesella cincta*. Genome Biology and Evolution, 8(7), 2106–2117.

25. Faddeeva-Vakhrusheva, A., et al. (2017). Coping with living in the soil: The genome of the parthenogenetic springtail *Folsomia candida*. BMC Genomics, 18, 493.

26. Fajana, H. O., Gainer, A., Jegede, O. O., Awuah, K. F., Princz, J. I., Owojori, O. J., & Siciliano, S. D. (2019). *Oppia nitens* CL Koch, 1836 (Acari: Oribatida): Current status of its bionomics and relevance as a model invertebrate in soil ecotoxicology. Environmental Toxicology and Chemistry, 38(12), 2593-2613.

27. Fajana, H. O., Jegede, O. O., James, K., Hogan, N. S., & Siciliano, S. D. (2020). Uptake, toxicity, and maternal transfer of cadmium in the oribatid soil mite, *Oppia nitens*: Implication in the risk assessment of cadmium to soil invertebrates. Environmental Pollution, 259, 113912.

28. Fajana, H. O., Hogan, N. S., & Siciliano, S. D. (2021). Does habitat quality matter to soil invertebrates in metal-contaminated soils? Journal of Hazardous Materials, 409, 124969.

29. Fajana, H. O., Rozka, T., Jegede, O., Stewart, K., & Siciliano, S. D. (2023). More than just a substrate for mites: Moss-dominated biological soil crust protected the population of the oribatid mite, *Oppia nitens*, against cadmium toxicity in soil. Science of the Total Environment, 857, 159553.

30. Fang, W. X., Dong, F. Y., Sun, E. T., Tao, D. D., Wang, Y., Xu, J. Y., … & Ye, C. J. (2020). De novo sequence of the mitochondrial genome of Tyrophagus putrescentiae (Acari: Sarcoptiformes) including 22 tRNA sequences and the largest non-coding region. Experimental and Applied Acarology, 80, 521–530.

31. Feyereisen, R. (2012). Insect CYP genes and P450 enzymes. In Insect molecular biology and biochemistry (pp. 236-316). Academic Press.

32. Gergócs, V., & Hufnagel, L. (2009). Application of oribatid mites as indicators. Applied Ecology and Environmental Research, 7(1), 79–98.

33. Giovannoni, S. J., Cameron Thrash, J., & Temperton, B. (2014). Implications of streamlining theory for microbial ecology. The ISME Journal, 8(8), 1553–1565.

34. Götz, S., Garcia-Gomez, J. M., Terol, J., Williams, T. D., Nagaraj, S. H., Nueda, M. J., Robles, M., Talon, M., Dopazo, J., & Conesa, A. (2008). High-throughput functional annotation and data mining with the Blast2GO suite. Nucleic Acids Research, 36(10), 3420–3435.

35. Gergócs, V., & Hufnagel, L. (2009). Application of oribatid mites as indicators. Applied ecology and environmental research, 7(1), 79–98.

36. Gregory, T. R. (2002). Genome size and developmental complexity. Genetica, 115, 131–146.

37. Gregory, T. R. (2005). Genome size evolution in animals. In The evolution of the genome (pp. 3-87). Academic Press.

38. Grbić, M., Van Leeuwen, T., Clark, R. M., Rombauts, S., Rouzé, P., Grbić, V., Osborne, E. J., Dermauw, W., Thi Ngoc, P. C., Ortego, F., & Hernández-Crespo, P. (2011). The genome of *Tetranychus urticae* reveals herbivorous pest adaptations. Nature, 479(7374), 487–492.

39. Gulia-Nuss, M., Nuss, A. B., Meyer, J. M., Sonenshine, D. E., Roe, R. M., Waterhouse, R. M., Sattelle, D. B., De La Fuente, J., Ribeiro, J. M., Megy, K., & Thimmapuram, J. (2016). Genomic insights into the *Ixodes scapularis* tick vector of Lyme disease. Nature Communications, 7(1), 10507.

40. Hamilton, W. L., Claessens, A., Otto, T. D., Kekre, M., Fairhurst, R. M., Rayner, J. C., & Kwiatkowski, D. (2017). Extreme mutation bias and high AT content in Plasmodium falciparum. Nucleic acids research, 45(4), 1889–1901.

41. Hoy, M. A., Waterhouse, R. M., Wu, K., Estep, A. S., Ioannidis, P., Palmer, W. J., Pomerantz, A. F., Simao, F. A., Thomas, J., Jiggins, F. M., & Murphy, T. D. (2016). Genome sequencing of the phytoseiid predatory mite *Metaseiulus occidentalis* reveals completely atomized Hox genes and superdynamic intron evolution. Genome Biology and Evolution, 8(6), 1762–1775.

42. Huot, J. L., Enkler, L., Megel, C., Karim, L., Laporte, D., Becker, H. D., Duchêne, A. M., Sissler, M., & Maréchal-Drouard, L. (2014). Idiosyncrasies in decoding mitochondrial genomes. Biochimie, 100, 95–106.

43. ISO 23266. (2020). Soil quality — Test for measuring the inhibition of reproduction in oribatid mites (*Oppia nitens*) exposed to contaminants in soil.

44. Jain, D., & Khurana, J. P. (2018). Role of pathogenesis-related (PR) proteins in plant defense mechanism. Molecular Aspects of Plant-Pathogen Interaction, 265–281.

45. Jalmi, S. K. (2022). The role of ABC transporters in metal transport in plants. In Plant Metal and Metalloid Transporters (pp. 55–71). Springer Nature Singapore.

46. Jegede, O. O., Hale, B. A., & Siciliano, S. D. (2019a). Multigenerational exposure of populations of *Oppia nitens* to zinc under pulse and continuous exposure scenarios. Environmental Toxicology and Chemistry, 38(4), 896–904.

47. Jegede, O. O., Awuah, K. F., Fajana, H. O., Owojori, O. J., Hale, B. A., & Siciliano, S. D. (2019b). The forgotten role of toxicodynamics: How habitat quality alters the mite, *Oppia nitens*, susceptibility to zinc, independent of toxicokinetics. Chemosphere, 227, 444–454.

48. Jeyaprakash, A., & Hoy, M. A. (2009). The nuclear genome of the phytoseiid *Metaseiulus occidentalis* (Acari: Phytoseiidae) is among the smallest known in arthropods. Experimental and Applied Acarology, 47, 263-273.

49. Jones, C. G., Lawton, J. H., & Shachak, M. (1994). Organisms as ecosystem engineers. In Ecosystem Management (pp. 130-147). Springer, New York, NY, USA.

50. Kanehisa, M., Sato, Y., & Morishima, K. (2016). BlastKOALA and GhostKOALA: KEGG tools for functional characterization of genome and metagenome sequences. Journal of Molecular Biology, 428(4), 726–731.

51. Keshavarz-Jamshidian, M., Verweij, R. A., Van Gestel, C. A., & Van Straalen, N. M. (2017). Toxicokinetics and time-variable toxicity of cadmium in *Oppia nitens* Koch (Acari: Oribatida). Environmental Toxicology and Chemistry, 36(2), 408-413.

52. Kumagai, Y., & Sumi, D. (2007). Arsenic: Signal transduction, transcription factor, and biotransformation involved in cellular response and toxicity. Annual Review of Pharmacology and Toxicology, 47(1), 243–262.

53. Laity, J. H., Lee, B. M., & Wright, P. E. (2001). Zinc finger proteins: New insights into structural and functional diversity. Current Opinion in Structural Biology, 11(1), 39–46.

54. Lavelle, P., Aubert, M., Blouin, M., Bureau, F., Mora, P., Margerie, P., Rossi, J. P., Barot, S., & Decaëns, T. (2006). Soil invertebrates and ecosystem services. European Journal of Soil Biology, 42, 3–15.

55. Li, Z., & Srivastava, P. (2003). Heat-shock proteins. Current Protocols in Immunology, 58(1), A-1T.

56. Lin, X., Wang, W., Ma, J., Sun, Z., Hou, H., & Zhao, L. (2021). Study on molecular level toxicity of Sb (V) to soil springtails: Using a combination of transcriptomics and metabolomics. Science of the Total Environment, 761, 144097.

57. Ludwig, M., Kratzmann, M., & Alberti, G. (1992). Observations on the proventricular glands (‘organes racémiformes’) of the oribatid mite Chamobates borealis (Acari, Oribatida): an organ of interest for studies on adaptation of animals to acid soils. Experimental & applied acarology, 15, 49–57.

58. Lynch, M., & Walsh, B. (2007). The origins of genome architecture (Vol. 98). Sunderland, MA: Sinauer Associates.

59. Meng, G., Li, Y., Yang, C., & Liu, S. (2019). MitoZ: A toolkit for animal mitochondrial genome assembly, annotation and visualization. Nucleic Acids Research, 47(11), e63–e63.

60. Mofiz, E., Seemann, T., Bahlo, M., Holt, D., Currie, B. J., Fischer, K., & Papenfuss, A. T. (2016). Mitochondrial genome sequence of the scabies mite provides insight into the genetic diversity of individual scabies infections. PLoS Neglected Tropical Diseases, 10(2), e0004384.

61. Norton, R. A. (1994). Evolutionary aspects of oribatid mite life histories and consequences for the origin of the Astigmata. In Mites: Ecological and Evolutionary Analyses of Life-History Patterns (pp. 99–135). Springer US.

62. Nota, B., Bosse, M., Ylstra, B., van Straalen, N. M., & Roelofs, D. (2009). Transcriptomics reveals extensive inducible biotransformation in the soil-dwelling invertebrate *Folsomia candida* exposed to phenanthrene. BMC Genomics, 10, 1–13.

63. Ohkubo, N. (2001). A revision of oppiidae and its allies (Acarina: Oribatida) of Japan 1. Genus Lasiobelba. Journal of the Acarological Society of Japan, 10, 97–109.

64. Owojori, O. J., & Siciliano, S. D. (2012). Accumulation and toxicity of metals (copper, zinc, cadmium, and lead) and organic compounds (geraniol and benzo[a]pyrene) in the oribatid mite *Oppia nitens*. Environmental Toxicology and Chemistry, 31(7), 1639–1648.

65. Owojori, O. J., Ademosu, O. T., Jegede, O. O., Fajana, H. O., Kehinde, T. O., & Badejo, M. A. (2019). Tropical oribatid mites in soil toxicity testing: Optimization of test protocol and the effect of two model chemicals (cadmium and dimethoate) on *Muliercula inexpectata*. Chemosphere, 218, 948–954.

66. Princz, J. I., Behan-Pelletier, V. M., Scroggins, R. P., & Siciliano, S. D. (2010). Oribatid mites in soil toxicity testing—The use of *Oppia nitens* (CL Koch) as a new test species. Environmental Toxicology and Chemistry, 29(4), 971–979.

67. Princz, J., Jatar, M., Lemieux, H., & Scroggins, R. (2018). Perfluorooctane sulfonate in surface soils: Effects on reproduction in the collembolan, *Folsomia candida*, and the oribatid mite, *Oppia nitens*. Chemosphere, 208, 757–763.

68. Salinas-Giegé, T., Giegé, R., & Giegé, P. (2015). tRNA biology in mitochondria. International Journal of Molecular Sciences, 16(3), 4518–4559.

69. Schäffer, S., Koblmüller, S., Klymiuk, I., & Thallinger, G. G. (2018). The mitochondrial genome of the oribatid mite *Paraleius leontonychus*: New insights into tRNA evolution and phylogenetic relationships in acariform mites. Scientific Reports, 8(1), 7558.

70. Schwager, E. E., Sharma, P. P., Clarke, T., Leite, D. J., Wierschin, T., Pechmann, M., Akiyama-Oda, Y., Esposito, L., Bechsgaard, J., Bilde, T., & Buffry, A. D. (2017). The house spider genome reveals an ancient whole-genome duplication during arachnid evolution. BMC Biology, 15, 1–27.

71. Schneider, A. (2011). Mitochondrial tRNA import and its consequences for mitochondrial translation. Annual Review of Biochemistry, 80(1), 1033–1053.

72. Sengupta, A., Bandyopadhyay, A., Sarkar, D., Hendry, J. I., Schubert, M. G., Liu, D., … & Pakrasi, H. B. (2024). Genome streamlining to improve performance of a fast-growing cyanobacterium Synechococcus elongatus UTEX 2973. Mbio, 15(3), e03530–23.

73. Seniczak, S., & Stefaniak, O. (1978). Microflora of the alimentary canal of *Oppia nitens* (Acarina, Oribatei). Pedobiologia, 18, 110–119.

74. Shao, Y., Wang, X. B., Zhang, J. J., Li, M. L., Wu, S. S., Ma, X. Y., Wang, X., Zhao, H. F., Li, Y., Zhu, H. H., & Irwin, D. M. (2020). Genome and single-cell RNA-sequencing of the earthworm *Eisenia andrei* identifies cellular mechanisms underlying regeneration. Nature Communications, 11(1), 1–15.

75. Šmarda, P., Bureš, P., Horová, L., Leitch, I. J., Mucina, L., Pacini, E., Tichý, L., Grulich, V., & Rotreklová, O. (2014). Ecological and evolutionary significance of genomic GC content diversity in monocots. Proceedings of the National Academy of Sciences, 111(39), E4096–E4102.

76. Spurgeon, D. J., Morgan, A. J., & Kille, P. (2008). Current research in soil invertebrate ecotoxicogenomics. Advances in Experimental Biology, 2, 133–326.

77. STAR: Dobin, A., Davis, C. A., Schlesinger, F., Drenkow, J., Zaleski, C., Jha, S., Batut, P., Chaisson, M., & Gingeras, T. R. (2013). STAR. Bioinformatics, 29(1), 15–21.

78. Stefaniak, O., & Seniczak, S. (1981). The effect of fungal diet on the development of *Oppia nitens* (Acari, Oribatei) and on the microflora of its alimentary tract. Pedobiologia, 21, 202–210.

79. Stelzer, C. P., Pichler, M., & Hatheuer, A. (2021). Linking genome size variation to population phenotypic variation within the rotifer, Brachionus asplanchnoidis. Communications Biology, 4(1), 596.

80. Suzuki, S., Kakuta, M., Ishida, T., & Akiyama, Y. (2014). GHOSTX: An improved sequence homology search algorithm using a query suffix array and a database suffix array. PloS One, 9(8), e103833.

81. van Straalen, N. M., & Feder, M. E. (2012). Ecological and evolutionary functional genomics: How can it contribute to the risk assessment of chemicals? Environmental Science & Technology, 46(1), 3–9.

82. Videvall, E. (2018). Plasmodium parasites of birds have the most AT-rich genes of eukaryotes. Microbial genomics, 4(2), e000150.

83. von Saltzwedel, H., Maraun, M., Scheu, S., & Schaefer, I. (2014). Evidence for frozen- niche variation in a cosmopolitan parthenogenetic soil mite species (Acari, Oribatida). PLoS One, 9(11), e113268.

84. Walker, B. J., Abeel, T., Shea, T., Priest, M., Abouelliel, A., Sakthikumar, S., Cuomo, C. A., Zeng, Q., Wortman, J., Young, S. K., & Earl, A. M. (2014). Pilon: An integrated tool for comprehensive microbial variant detection and genome assembly improvement. PloS One, 9(11), e112963.

85. Warren, J. M., & Sloan, D. B. (2021). Hopeful monsters: unintended sequencing of famously malformed mite mitochondrial tRNAs reveals widespread expression and processing of sense–antisense pairs. NAR Genomics and Bioinformatics, 3(1), lqaa111.

86. Wickings, K., & Grandy, A. S. (2011). The oribatid mite *Scheloribates moestus* (Acari: Oribatida) alters litter chemistry and nutrient cycling during decomposition. Soil Biology and Biochemistry, 43, 351–358.

87. WindowMasker: Morgulis, A., Gertz, E. M., Schäffer, A. A., & Agarwala, R. (2006). WindowMasker. Bioinformatics, 2, 134–141.

88. Wu, C., Jordan, M. D., Newcomb, R. D., Gemmell, N. J., Bank, S., Meusemann, K., … & Buckley, T. R. (2017). Analysis of the genome of the New Zealand giant collembolan (*Holacanthella duospinosa*) sheds light on hexapod evolution. BMC Genomics, 18, 1–19.

89. Wu, H., Zhang, Z., Hu, S., & Yu, J. (2012). On the molecular mechanism of GC content variation among eubacterial genomes. Biology Direct, 7, 1–16.

90. Xue, X. F., Deng, W., Qu, S. X., Hong, X. Y., & Shao, R. (2018). The mitochondrial genomes of sarcoptiform mites: are any transfer RNA genes really lost?. BMC genomics, 19, 1–11.

91. Yang, B., & Li, C. (2016). Characterization of the complete mitochondrial genome of the storage mite pest *Tyrophagus longior* (Gervais) (Acari: Acaridae) and comparative mitogenomic analysis of four acarid mites. Gene, 576(2), 807-819.

92. Zhan, X. B., Chen, B., Fang, Y., Dong, F. Y., Fang, W. X., Luo, Q., … & Sun, E. T. (2021). Mitochondrial analysis of oribatid mites provides insights into their atypical tRNA annotation, genome rearrangement and evolution. Parasites & Vectors, 14(1), 221.

93. Zhang, F., Ding, Y., Zhou, Q. S., Wu, J., Luo, A., & Zhu, C. D. (2019). A high-quality draft genome assembly of *Sinella curviseta*: A soil model organism (Collembola). Genome Biology and Evolution, 11(2), 521–530.

94. Zhu, Y., Wu, X. Y., Liu, Y. X., Zhang, J. Y., & Lin, D. H. (2020). Integration of transcriptomics and metabolomics reveals the responses of earthworms to the long-term exposure of TiO2 nanoparticles in soil. Science of the Total Environment, 719, 137492.

95. Zwarycz, A. S., Nossa, C. W., Putnam, N. H., & Ryan, J. F. (2016). Timing and scope of genomic expansion within Annelida: Evidence from homeoboxes in the genome of the earthworm *Eisenia fetida*. Genome Biology and Evolution, 8(1), 271–281.

